# GPR174 Antagonism: Structure, Function, and Dynamics

**DOI:** 10.1101/2025.09.18.677133

**Authors:** Vijay Kumar Bhardwaj, Alemayehu Gorfe

## Abstract

GPR174 is an immune-restricted G-protein–coupled receptor (GPCR) constitutively activated by lysophosphatidylserine (LysoPS). Elevated LysoPS in the tumor microenvironment may sustain GPR174 activity, promoting immunosuppression and resistance to cancer immunotherapies. Here, we modeled GPR174 bound to an antagonist mPS (modified LysoPS) and performed extensive molecular dynamics (MD) simulations in a heterogeneous lipid bilayer, with parallel simulations of the LysoPS-bound receptor for comparison. mPS binding inactivated GPR174 and resulted in reduced conformational dynamics, persistent hydrogen bonding interactions, and selective interactions with transmembrane helix 1. In contrast, LysoPS exhibited greater conformational flexibility, multiple binding poses, and transient acyl chain displacement into the membrane. Network analysis revealed that LysoPS engaged conserved activation motifs (PIF, DRY, N/DPxxY) to couple the ligand binding site to the G-protein interface, whereas these pathways were disrupted by mPS. Protein–lipid analyses further suggested that membrane lipids, including phosphatidylinositol (PIP2), modulate ligand dynamics and receptor conformational states. Collectively, these findings highlight distinct ligand-specific mechanisms of GPR174 modulation and provide a framework for rational design of selective antagonists with immunotherapeutic potential.

## Introduction

G protein-coupled receptors (GPCRs) are one of the most prominent protein families encoded by the human genome. These transmembrane proteins are responsible for channeling extracellular signals into the cells to regulate many physiological functions^1,2^. GPCRs represent a historically significant and highly successful class of pharmacological targets; approximately 35% of currently approved drugs directly target GPCRs, representing roughly 12% of all protein targets for approved therapeutics^3^. This presents an opportunity to investigate the therapeutic potential of other, less-characterized members of this large protein superfamily.

One such GPCR is GPR174, also known as FKSG79 and GPCR17 or LPS3, that belongs to the P2Y receptor family of orphan GPCRs^4–6^. GPR174 is highly expressed in a relatively small number of tissues, including lymph nodes, spleen, thymus, and bone marrow. Within the lymphatic system, GPR174 is prominently expressed in T and B cells, especially in regulatory T cells (Tregs)^7^. In recent studies, the activation of GPR174 by lysophosphatidylserine (LysoPS) was shown to negatively regulate the function of both T and B cells by increasing the production of intracellular cyclic adenosine monophosphate (cAMP)^6–8^. Given the established role of regulatory Tregs in suppressing excessive inflammatory responses, inhibiting GPR174 could represent a potential therapeutic strategy for autoimmune disorders. Genetic studies have identified single-nucleotide polymorphisms (SNPs) in the GPR174 gene that are associated with autoimmune disorders such as Grave’s disease^4^ and Addison’s disease^9^. Additionally, altered expression levels of GPR174 have been linked to the occurrence of vasovagal syncope^10^. Interestingly, in synthetic liposome, phosphatidylserine (PS) was found to be five times more potent than LysoPS in stimulating GPR174-dependent cAMP production^11^. Under normal conditions, PS is mainly confined to the inner leaflet of the plasma membrane. However, PS is translocated to the outer leaflet by phospholipid scramblases during certain physiological processes like stress and apoptosis or during the activation of various cell types, including different immune cells or tumor cells^12–14^. Hence, high concentrations of PS are considered a major source of tumor-mediated immunosuppression and may play a role in resistance to cancer immunotherapies^15^. Thus, compounds that can modulate GPR174 activity, especially GPR174 antagonists, might be useful both in biological research and as candidate drugs for the control of various types of cancers and autoimmune diseases.

A recently published study identified compound 7d (hereafter mPS) as a potent GPR174 antagonist^16^. Structurally, mPS is derived from the agonist LysoPS via modifications including acetylation of the (S)-amino group, conversion of the glycerol-fatty acid ester linkage to an amide bond, and the use of a meta-substituted benzene system as a non-fatty acid surrogate^16^. The chemical structures of LysoPS and mPS are shown in Fig. 1a-b. However, the precise molecular mechanism underlying GPR174 inhibition by mPS remains to be elucidated. The publication of the cryo-EM structure of the GPR174-LysoPS-Gs complex^17–19^ now provides a crucial structural basis for investigating the mechanisms of receptor activation and inhibition, as well as for identifying novel small-molecule inhibitors.

**Fig. 1.**
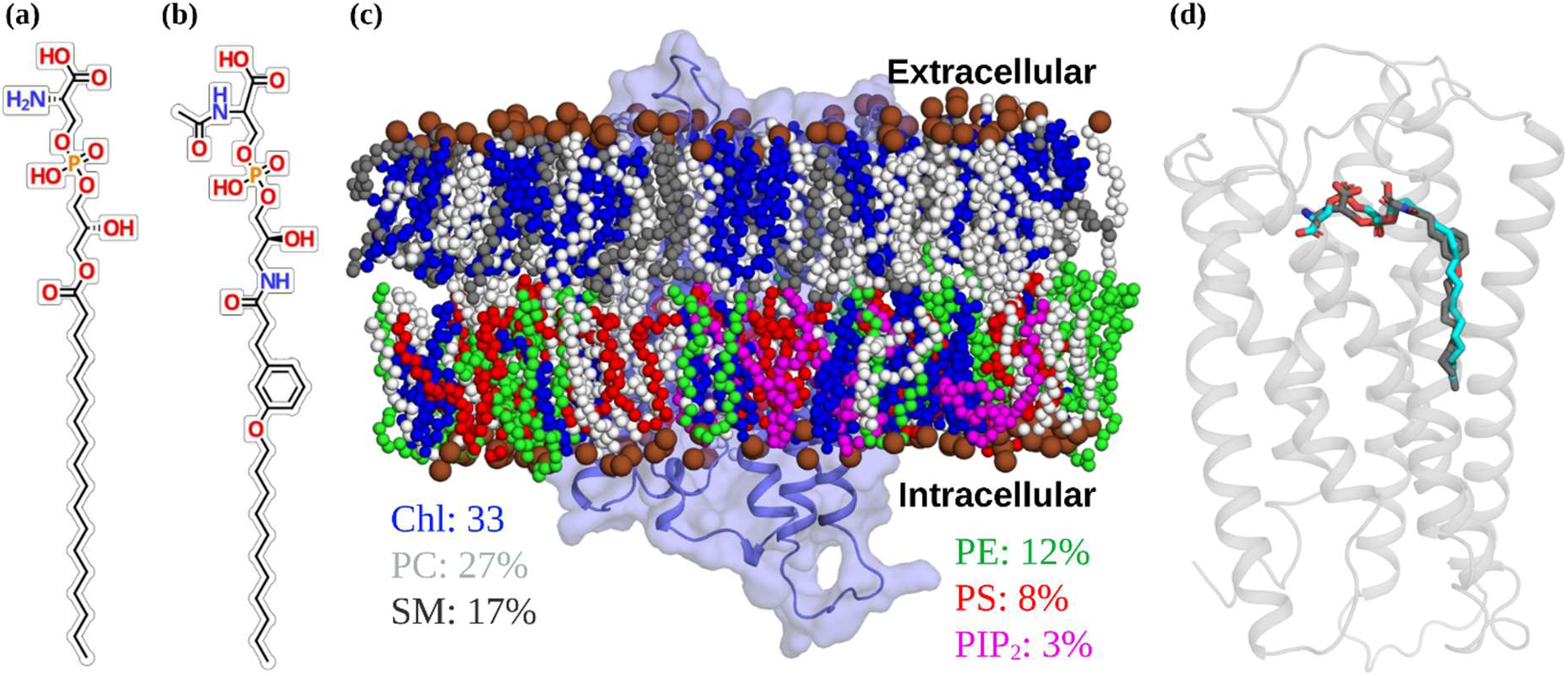
Structural representation of GPR174, its lipid environment, and ligand binding. Chemical structure of **(a)** LysoPS and **(b)** mPS. **(c)** GPR174 embedded in a lipid bilayer representing a realistic plasma membrane composition. The acyl chains of lipid species are color-coded and labeled with their respective molar ratios of total lipids. All P atoms are shown as brown spheres. **(d)** A comparison of the ligand binding poses between LysoPS (experimental) and mPS (docked). Chl: Cholesterol; PC: Phosphatidylcholine; PE: Phosphatidylethanolamine; PlaPE: Plasmalogen PE; PIP2: Phosphatidylinositol; PS: phosphatidylserine; SM: sphingomyelin.

This study aims to elucidate the molecular mechanism of GPR174 inhibition by mPS. Using modeling and long-timescale molecular dynamics (MD) simulations, we compare the binding and conformational effects of mPS with the endogenous agonist LysoPS, examine their influence on conserved activation motifs and signaling pathways, and evaluate the role of membrane lipids in modulating receptor dynamics. These resulting insights provide a framework for the rational design of selective GPR174 antagonists with potential therapeutic relevance.

## Results

### Modeling of GPR174 complexes

While the experimental structure of GPR174 in its active state bound to the endogenous agonist LysoPS has been recently determined^17–19^, structural information regarding antagonist-bound conformations remains unavailable. Given a recent report demonstrating the conversion of LysoPS to an antagonist upon acylation of its L-serine moiety^16^, we employed computational docking to model the binding of the potent antagonist 7d (referred to as mPS in this study) within the GPR174 ligand-binding pocket (Fig. 1d, Fig. S1-S2). To ensure the reliability of our docking protocol, we performed re-docking of LysoPS into the same binding site and compared the resulting pose with the experimentally determined structure. The conservation of all key interactions and a low root-mean-square deviation (RMSD) of 0.42 Å from the experimental structure support the reliability of our docking methodology. Comparative analysis of the top-ranked mPS docking pose with the experimental LysoPS structure revealed a potentially distinct binding mode characterized by the loss of interactions with R75^2.60^, Y99^3.33^, and F169^ECL2^, and the addition of novel interactions with R18^1.31^ and Y22^1.35^. These newly formed interactions with mPS, mediated by its added acyl group, involved residues located on TM1, which did not interact with LysoPS in the active state structure. However, it should be noted that mPS was docked on the rigid active-state receptor and does not truly represent an inactive conformation. Based on these observations, we hypothesized that the differential binding mode observed for mPS may drive conformational changes towards an inactive conformation of GPR174. To test this hypothesis and assess the impact of mPS binding on the overall receptor dynamics, we conducted microsecond timescale MD simulations.

### Structural stability of protein-ligand-membrane complexes

The experimentally determined GPR174-LysoPS structure and the top-ranked docked pose of the GPR174-mPS complex were embedded in a heterogeneous lipid bilayer to mimic the plasma membrane environment surrounding GPR174. Both systems were subjected to 2 µs of MD simulations, performed in duplicates. To assess the overall structural stability of systems during the simulations, we monitored RMSD of the Cα atoms (Fig. 2a) and membrane thickness quantified as the average phosphorus-to-phosphorus (P-P) distance across the bilayer (Fig. 2b).

**Fig. 2.**
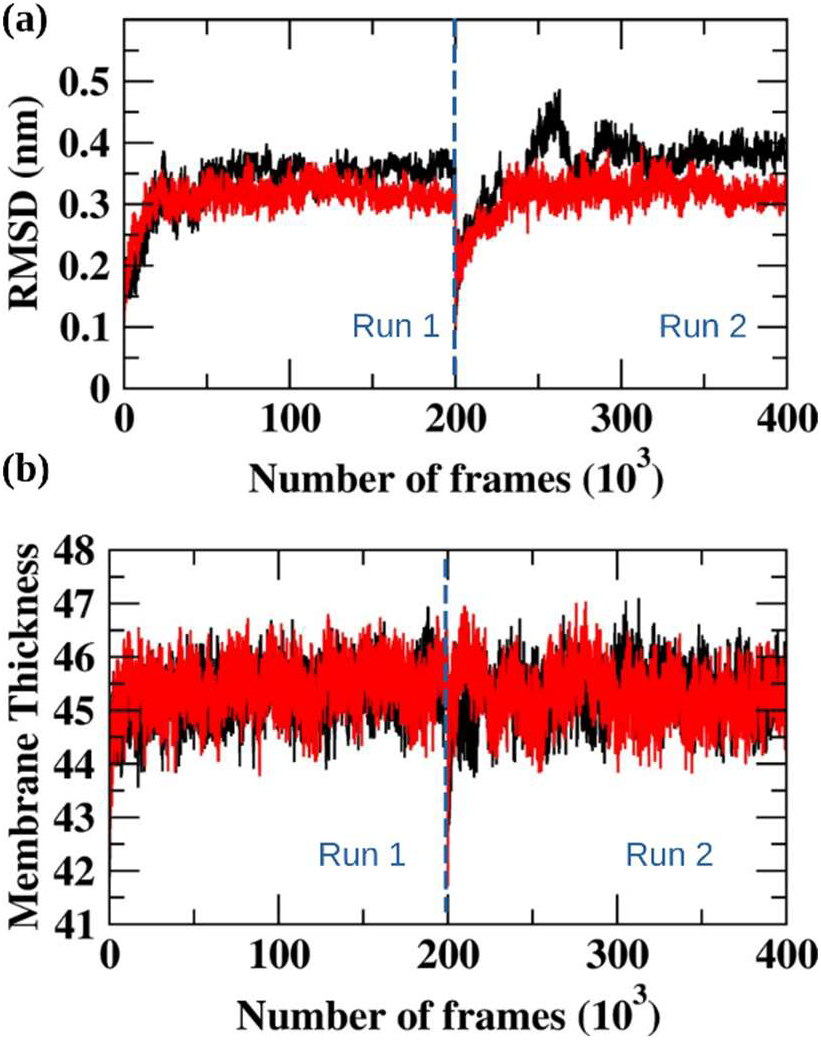
Overall structural stability of simulated systems. **(a)** Time evolution of protein Cα atoms RMSD and **(b)** membrane thickness for LysoPS (black) and mPS (red) systems. Data are shown for two concatenated trajectories separated by a dashed blue line, indicating Run 1 and Run 2.

In each system and simulation, equilibration was achieved within ∼250 ns of the simulations (Gig. 2). The mPS-bound GPR174 consistently displayed a slightly lower RMSD than the LysoPS-bound GPR174, suggesting a trend towards better conformational stability in the presence of the antagonist. The P-P distance (Fig. 2b) revealed a stable lipid bilayer environment for both systems with an average value of 45.3 ± 0.4 Å. A similar P-P distance was observed in a study that compared 18 biomembrane systems with lipid compositions corresponding to eukaryotic, bacterial, and archaebacterial membranes^20^.

In addition, residue-level flexibility was assessed by root-mean-square fluctuation (RMSF) analysis (Fig. S3). Both complexes exhibited similar fluctuation profiles across most transmembrane helices, whereas enhanced flexibility was observed in the intracellular loop regions (ICL1 and ICL2) of the LysoPS-bound system compared to the mPS-bound system.

### mPS binding drives GPR174 to an inactive state

The molecular mechanisms underlying GPCR activation governed by allosteric processes are extensively studied, linking agonist binding to the characteristic outward displacement of TM6^21–23^. Since the antagonist mPS was docked on the already active state of GPR174, we anticipated mPS binding to transform it to its inactive form. In order to capture the deactivation process, specifically focusing on the movement of TM6 relative to TM3 and TM2 and monitored the angle formed between vectors representing these transmembrane helices (Fig. 3a,b, Fig. S4a-b). LysoPS and mPS bound structures had similar angles between TM6-TM3 (∼38°) and TM6-TM2 (∼64°) at the start of simulations (Fig. S4a-b). The mPS-bound system maintained a consistently smaller TM6-TM3 angle (22-32°) while the LysoPS system fluctuated between 35° and 55°. Similarly, the mPS-bound system displayed a consistent trend towards smaller TM6-TM2 angles, generally ranging from 40° to 55° throughout the simulation. The LysoPS system showed greater variability in the TM6-TM2 angle, fluctuating between approximately 55° and 80°. To complement these results, we calculated the distance between the cytosolic sides of TM6 and TM3 (Fig. 3c, Fig. S4c). As expected, the mPS system exhibited a smaller distance (around 8-10 Å) than the LysoPS system, which showed a wider distribution with distances predominantly in the range of 11-17 Å. Collectively, these simulations reveal that mPS binding stabilizes TM6 in a conformation characterized by reduced TM6-TM3 and TM6-TM2 angles, and a decreased distance between the cytosolic ends of TM6 and TM3, all indicative of the inactive state of GPR174. Conversely, LysoPS binding allows for a more dynamic TM6, potentially reflecting active or intermediate states.

**Fig. 3.**
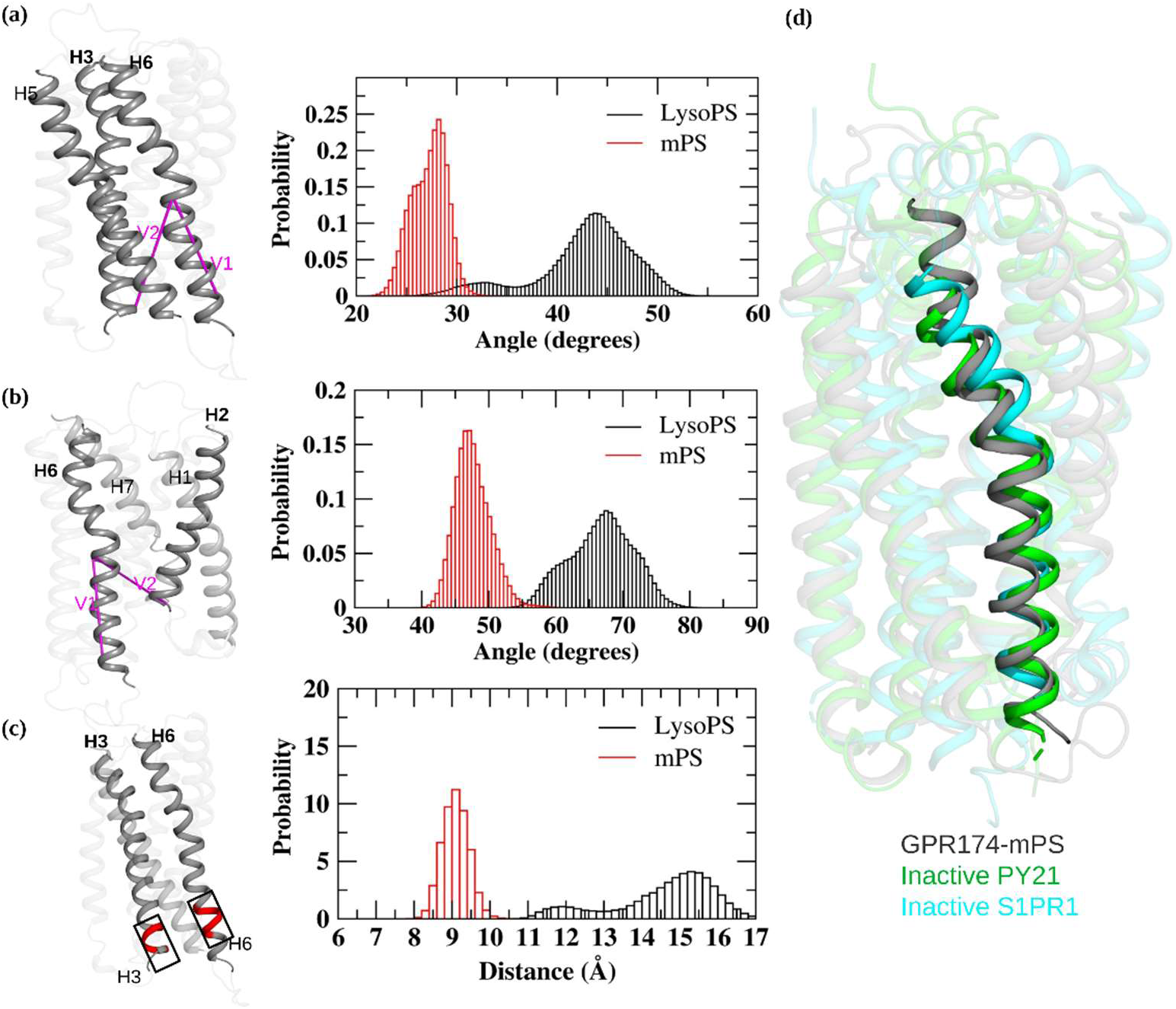
mPS binding inactivates GPR174. Probability distributions of interhelical angles between **(a)** TM6-TM3 and **(b)** TM6-TM2 for LysoPS-bound (black) and mPS-bound (red) GPR174. Angles were computed between vectors (V1 and V2) as shown in the cartoon models to the left. **(c)** Distribution of the distance between TM3 and TM6 at the cytosolic end, measured as the center-of-mass distance between the Cα atoms of residues highlighted in red in the cartoon representation. **(d)** Superposition of the last frame of Run1 from the mPS-bound GPR174 simulation with experimentally solved inactive GPCR structures: P2Y1 (PDB:4XNW), S1PR1 (PDB:3V2Y).

We performed a comparative structural analysis to establish that the observed reduction in angles and distance in the mPS-bound simulation corresponds to an inactive conformation of GPR174. To this end, the last frame of one of the mPS-bound simulations was superimposed onto experimentally determined structures of antagonist-bound GPCRs, including P2Y1, which represents the most closely related receptor with an experimentally solved inactive state structure (Fig. 3d). TM6 of the mPS-bound GPR174 showed close alignment with the TM6 helices from the inactive P2Y1 and S1PR1 structures, particularly towards the cytosolic end. The compelling alignment of the GPR174-mPS TM6 with that of the inactive P2Y1 and S1PR1 structures provided strong evidence that the simulations captured an inactive conformation of GPR174.

### Comparison of active and inactive GPR174 ensembles

Understanding the activation and deactivation processes of target proteins is essential for developing effective therapeutic strategies. Our analysis of GPR174 in both active and inactive states provides a framework for elucidating the structural network and dynamics. To this end, we employed PSNtools^24^ to compare the shortest communication paths in the equilibrated portions of the trajectories of GPR174 bound to LysoPS and mPS (Fig. 4a-b). On the extracellular end, the agonist LysoPS engaged with a distinct set of residues compared to the antagonist mPS.

**Fig. 4.**
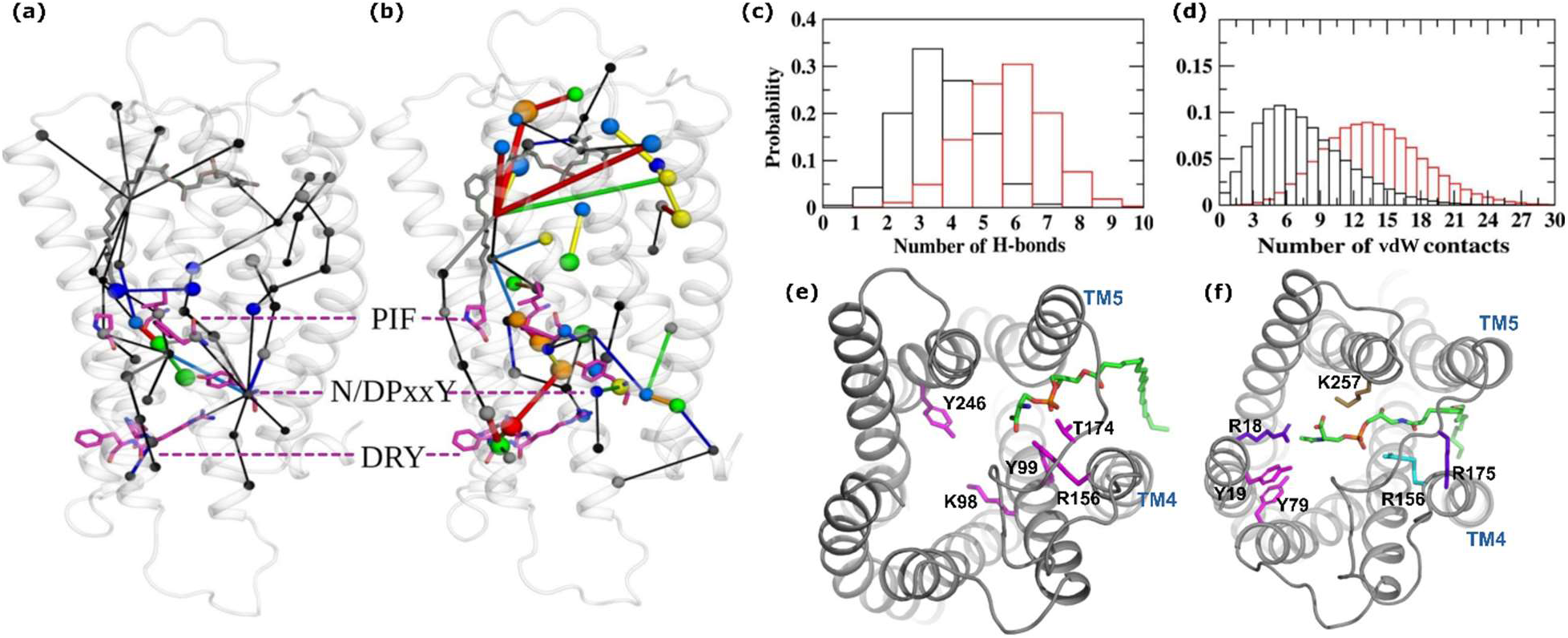
Analysis of intramolecular communication pathways and ligand binding. Network diagrams derived from MD simulations showing shortest communication pathways in **(a)** LysoPS-bound and **(b)** mPS-bound GPR174. Nodes represent residues, and edges represent interactions. Thickness and color represent strength and frequency (Red>Orange>Yellow>Green>Blue>Black) of interactions, respectively. Dashed magenta lines indicate the relative positions of conserved motifs PIF, N/DPxxY, and DRY. Distributions of **(c)** number of hydrogen bonds and **(d)** number of van der Waals contacts formed by LysoPS (black) and mPS (red) with GPR174. Structural mapping of representative hydrogen bond interactions in **(e)** LysoPS-bound and **(f)** mPS-bound GPR174 complexes. Interacting residues are color-coded according to hydrogen bond frequencies: 20-30% (magenta), 30-40% (cyan), 40-50% (brown), >50% (purple-blue). TM4 and TM5 are labeled for spatial reference.

Specifically, LysoPS interactions were primarily localized to K257^6.62^, L179^5.32^, V170^ECL2^ and R175^ECL2^. While mPS also interacts with F254^6.59^, it exhibited a higher preference for residues R18^1.31^, Y22^1.35^, F152^4.60^, and R156^4.64^. These interactions led to distinct signaling pathways, with TM4 forming connections to TM3 and ECL2, while TM1 was linked to TM7 and TM5 on the extracellular end. Clearly, LysoPS and mPS are characterized by different communication pathways on the extracellular part of GPR174. However, none of these pathways directly extended into the transmembrane domain.

Conserved motifs PIF, N/DPxxY and DRY on GPCRs were previously shown to play key roles in relaying signal through the transmembrane region, thereby coupling ligand binding to G protein activation ^25,26^. The PIF motif having residues on three TM helices acts as a micro-switch that relays signal from the extracellular part to the cytoplasmic end of GPCRs. LysoPS binding resulted in a communication pathway engaging the PIF motif, where G193^5.46^ bridges the interaction with P197^5.50^ and I106^3.40^. The signal from P197^5.50^ and I106^3.40^ converges at L109^3.43^, where it bifurcates into two distinct pathways: one leading to Y292^7.52^ of the N/DPxxY motif and the other to the N-terminus of ICL3. Y292^7.52^ subsequently transmits the signal to the C-terminus of ICL3, a region known to interact with the G protein, thereby facilitating downstream signaling. Furthermore, Y292^7.52^ acts as a key node, initiating communication pathways establishing connections between TM7 and both TM1 and TM2, as well as between TM2 and TM3 (SI movie 1). In contrast to LysoPS, analysis of the mPS-bound ensemble revealed an absence of direct communication pathways linking the ligand-binding site to the PIF motif. Additionally, no direct connections were observed between the three conserved motifs. None of the signals traveled to either end of the ICL3, and communication pathways arising from Y292^7.52^ of the conserved DRY motif were lost (SI movie 2).

### LysoPS and mPS interact with a different set of residues in the binding pocket

We analyzed the differences in hydrogen bonding and vdW interactions between LysoPS and mPS to understand how each ligand differentially modulates GPR174 activation and inhibition. The distribution of hydrogen bonds (Fig. 4c) revealed a clear contrast in interaction profiles. mPS formed a higher and more consistent number of hydrogen bonds with GPR174, with most frames showing 5-8 hydrogen bonds (Fig. S5a). In contrast, LysoPS displayed a broader and more left-skewed distribution, with the majority of frames exhibiting only 2-4 hydrogen bonds, indicative of less persistent polar interactions. A similar trend was observed in the distribution of vdW contacts (Fig. 4d), which reflects hydrophobic engagement between the acyl chain of each ligand and the receptor. mPS consistently forms 30-50 contacts per frame (Fig. S5b), while LysoPS shows a broader distribution skewed toward lower values, typically between 15 and 30 contacts. The reduction in the number of contacts for LysoPS between frames ∼550 and ∼1500 corresponds to the ligand acyl chain partially moving out to interact with the membrane (see below). Also, the reduction in the number of hydrogen bonds for LysoPS aligns with network analysis (Fig. 4a) and clustering analyses (see below), which revealed substantial conformational diversity. Notably, the head group of LysoPS showed lateral displacement away from its starting position, shifting toward TM4-TM6. This movement likely disrupted stable hydrogen bonding by repositioning polar atoms away from key interacting residues, thereby reducing both the number and persistence of hydrogen bonds over the trajectory.

To further investigate these differences in interaction profiles, we analyzed the residue-level hydrogen bond occupancy for both ligands (Fig. 4e-f). LysoPS interacts with a wide set of residues on TM2, TM5, TM6, and ECL2, including Y246^6.51^ (24%), T174^ECL2^ (23%), Y99^3.33^ (22%), and E190^5.43^ (17%). While these interactions contribute to transient stabilization, their moderate occupancy reflects the flexible and shifting binding pose of the agonist (Fig. 4e). In contrast, mPS established a well-defined and persistent interaction network with GPR174. R175^ECL2^ exhibited the highest hydrogen bond occupancy at 74%. Notably, this interaction arose as a direct consequence of the chemical modification that converted LysoPS into mPS by the replacement of the glycerol-fatty acid ester linkage with an amide bond, introducing a carbonyl oxygen that forms a stable hydrogen bond with R175^ECL2^. Interestingly, mPS formed stable hydrogen bonds with R18^1.31^ (52% occupancy) and Y19^1.32^ (18% occupancy) on TM1 (Fig. 4f). These interactions were completely absent in the LysoPS-bound complex, highlighting a distinct binding pose for the antagonist and suggesting a potential role for TM1 in stabilizing the inactive conformation of GPR174.

### Analyzing the diversity of ligand conformations

Given the conformational dynamics indicated by the differences in network analysis and the number of ligand-protein contacts (Fig. 4), especially the variations observed for the ligand head moieties, we sought to determine if distinct and stable conformational states were populated during the simulations. Therefore, an RMSD-based clustering (0.18 nm cutoff) analysis was conducted on equilibrated trajectories to group structurally similar snapshots and extract the predominant conformations of LysoPS and mPS. This analysis revealed strikingly different binding orientations and sites for the agonist and the antagonist, highlighting distinct binding modes (Fig. 5a-b). For mPS, the clustering converged into a very limited number of conformations (14 total clusters), with a single cluster overwhelmingly dominating the ensemble, accounting for 94% (red) of the simulation frames (Fig. 5b). The representative structures of the minor clusters showed high similarity to this dominant pose, indicating that mPS adopts a highly stable and well-defined conformation when bound to GPR174. Conversely, LysoPS displayed significant conformational heterogeneity, partitioning into a large number (226) of distinct clusters. The population was broadly distributed across these clusters, with the three most populated accounting for only 19% (red), 17% (green), and 8% (blue), collectively representing less than half of the simulation time (Fig. 5a). Visual inspection of the representative structures for LysoPS confirmed substantial conformational diversity. Therefore, the clustering analysis demonstrated that while the antagonist mPS maintains a stable, dominant binding pose, the agonist LysoPS dynamically samples a diverse range of distinct conformations within the GPR174 binding pocket.

**Fig. 5.**
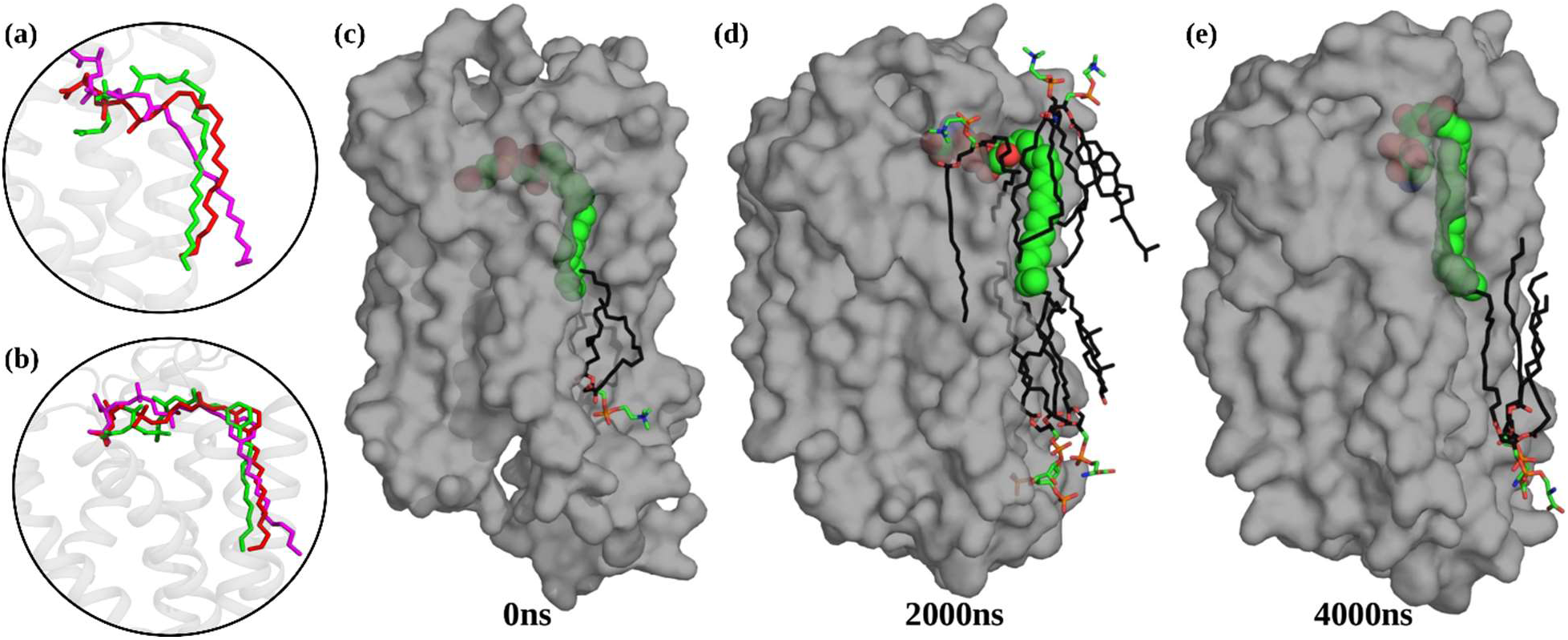
Analysis of LysoPS acyl chain flexibility and binding dynamics. The representative clustered conformations of **(a)** lysoPS and **(b)** mPS during simulations. **(c-e)** Snapshots of ligand conformations at 0ns, 2000ns, and 4000ns. The GPR174 receptor is shown in surface representation (gray), LysoPS as spheres (atom-colored), and membrane lipids within 4Å of LysoPS as stick model (acyl chain: black; head: atom-colored).

### Characterization of LysoPS acyl chain lateral dynamics

The analysis of representative structures from the most populated clusters revealed that the acyl chain of LysoPS transitioned from its initial binding site (hydrophobic pocket formed by helices 4 and 5) to interact with membrane lipids. This prompted us to further investigate the potential unbinding of LysoPS from its established binding site by extending one of the MD runs of LysoPS-bound GPR174 to 4µs. Quantification of vdW contacts between LysoPS and surrounding lipids demonstrated a dynamic interaction profile (Fig. S6). Initially, LysoPS exhibited a minimal number of contacts with membrane lipids. As the simulation progressed, the number of contacts between LysoPS and the bilayer lipids significantly increased (average ∼12 contacts between 1000ns and 3000ns), indicative of the acyl chain’s engagement with the membrane environment. However, towards the latter phase of the extended simulation, a decrease in interactions was observed, potentially reflecting a disengagement from the membrane. Structural analysis (Fig. 5d) suggested an intermediate level of membrane engagement (time 2000ns), thereafter, the acyl chain of LysoPS appeared to have re-entered the hydrophobic pocket at time 4000ns (Fig. 5e). These observations were further strengthened by the analysis of vdW contacts between LysoPS acyl chain and side chains of hydrophobic residues lining helices 4 and 5 (Fig. 5e). As expected, the number of contacts was initially high, reflecting engagement within the pocket, subsequently decreasing as the ligand transitioned out and re-engaging with the pocket towards the end of the simulation. To further validate this observation, we quantified the number of membrane lipid atoms occupying the hydrophobic cavity normally engaged by the LysoPS acyl chain (Fig. S6b). Between 1000 ns and 3000 ns, the number of lipid atoms within the pocket increased from ∼2 to ∼6, coinciding with the period of reduced ligand engagement. Toward the end of the simulation, lipid occupancy declined as the LysoPS tail re-entered the pocket. These findings suggest that membrane lipids may transiently occupy the cavity and potentially compete with or displace the LysoPS acyl chain from its binding site.

This complex behavior, characterized by both membrane interaction and re-entry into the hydrophobic pocket, suggested a dynamic equilibrium or a multi-step unbinding of LysoPS, rather than a simple reversible dissociation. The transient occupation of the binding pocket by lipids further points to an active role of the membrane in modulating ligand dynamics. These observations further motivated a detailed investigation into the role of membrane lipids in stabilizing distinct conformational states of the ligand-receptor complexes.

### Lipid-Protein Interactions

Membrane lipids serve as critical regulators of protein function, governing the activation and modulation of GPCRs, ion channels, and other membrane proteins^27–29^. We subjected the equilibrated trajectories of both LysoPS- and mPS-bound GPR174 to PyLipID tool^30^ for the identification and characterization of specific interactions and binding sites of cholesterol (Chl) and phosphatidylinositol (PIP2). This focus was prompted by the established roles of both lipids in the stabilization of different conformational states of GPCRs^31,32^. The occupancy profiles of identified lipid-binding sites are summarized in Fig. S7-S8. A binding site was defined as common if at least three residues matched between the two systems, allowing for a positional tolerance of ±1 residue. We identified five Chl binding sites on each of the LysoPS- and mPS-bound GPR174 structures, with two located on the cytoplasmic side and four towards the extracellular region (Fig. S9a-b). Notably, we captured a Chl binding site consistent with that observed in the experimental structure of LysoPS-bound GPR174 (Fig. S9c). However, this site was only present in the mPS-bound system. In the LysoPS system, the same pocket was occupied by a PIP2, likely preventing Chl from accessing the site. In contrast to Chl, PIP2 binding patterns were markedly different between the two systems. The LysoPS-bound GPR174 exhibited a single PIP2 binding site, whereas the mPS-bound system revealed three distinct PIP2 interaction sites (Fig. 6). None oft hese sites were conserved across the two systems, suggesting that PIP2 interactions are highly dependent on the ligand state of GPR174. Notably, in the LysoPS-bound structure, the PIP2 occupied the pocket between helices 4 and 5 throughout the simulation, consistent with our observation of LysoPS tail displacement by membrane lipids. Thus, PIP2 binding in the LysoPS system may transiently stabilize an intermediate state. Conversely, the presence of multiple PIP2 binding sites in the mPS-bound system may contribute to the stabilization of an inactive receptor conformation. Together, these results highlight a lipid-dependent mechanism of receptor modulation in which PIP2 plays distinct roles depending on the ligand state of GPR174.

**Fig. 6.**
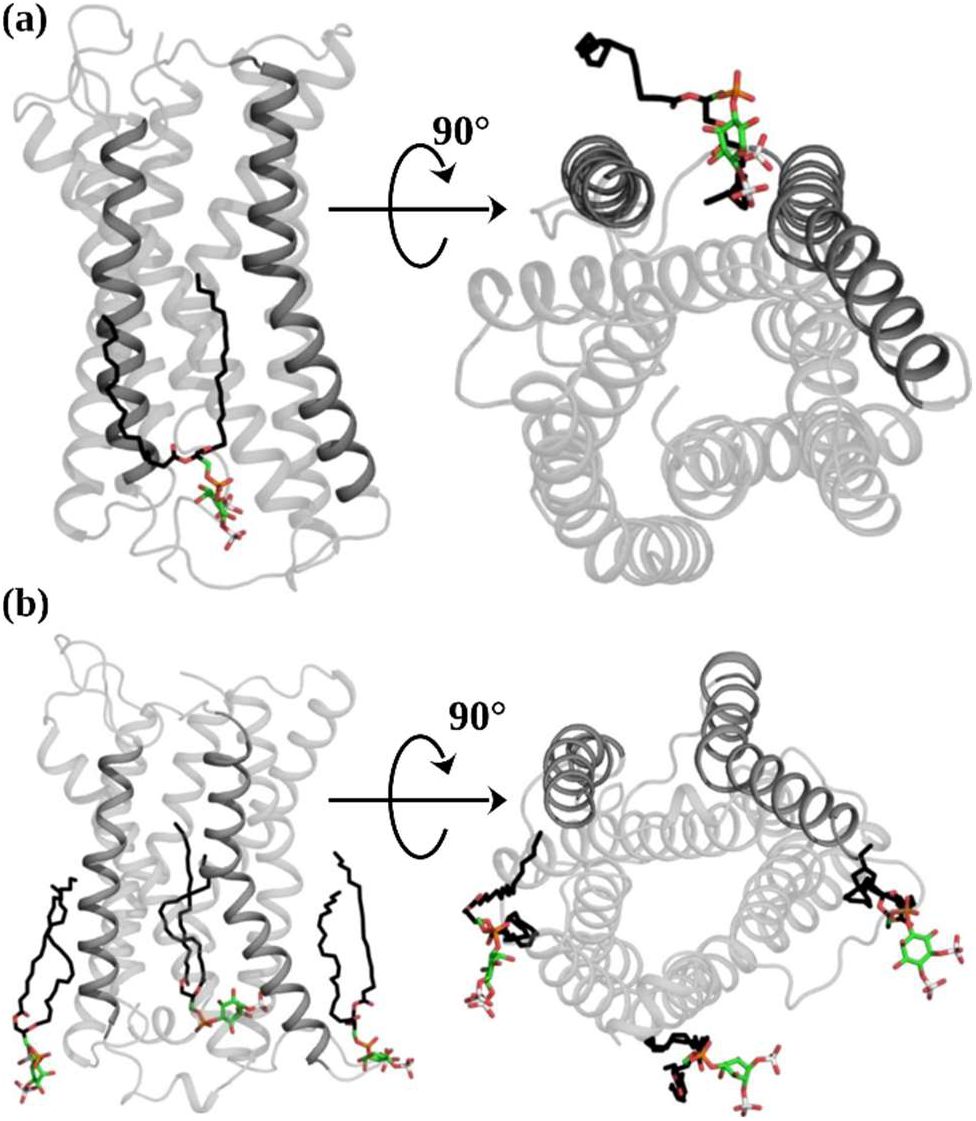
Protein-lipid interactions. Analysis of PIP2 binding sites for **(a)** LysoPS-bound and **(b)** mPS-bound GPR174. The acyl chains are colored black, while the head group of PIP2 is atom-colored in the stick model.

## Discussions

Recent structural analysis of GPR174 in its active state bound to LysoPS has provided valuable insights for understanding the mechanisms of its agonist-dependent activation^17–19^. However, these structures capture static conformations under defined experimental conditions, potentially lacking critical details about the conformational ensembles of GPCRs^33^. To bridge this gap, MD simulations have emerged as a powerful complement to experimental structural data, providing dynamic insights and serving as a key approach for investigating receptor mechanisms^34–38^.

Elucidating the molecular mechanisms of GPCR inhibition is essential for informing the rational design of antagonists. In the present study, we employed atomistic MD simulations of the GPR174 receptor to elucidate its deactivation mechanism. We used the experimentally determined active-state GPR174-LysoPS structure^17^ as a template for generating an initial GPR174-mPS structure through docking (Fig. 1d). Since the docked pose is unlikely to adequately represent the antagonist-bound conformation of GPR174, we performed MD simulations, anticipating that mPS binding would drive GPR174 towards an inactive conformation. All simulations were performed in a complex PM lipid environment, as prior research has demonstrated the critical role of lipid composition in modulating GPCR function and structural stability^39,40^. To evaluate the overall stability and convergence of the trajectories, we monitored RMSD (Fig. 2a) and membrane thickness (Fig. 2b) as an indicator of bilayer integrity. Next, we analyzed the angle formed by TM6 with TM3 and TM2 towards the cytoplasmic end in both LysoPS- and mPS-bound trajectories to characterize active and inactive conformations of GPR174 (Fig. 3a-b). To complement these results, we also calculated the distance between TM6 and TM3 (Fig. 3c). The outward displacement of TM6 is a characteristic of class A GPCR activation^21^ and is commonly used to distinguish between inactive and active receptor states^36,41,42^. Our analysis clearly showed a decrease in the angles formed by TM6 with selected helices, suggesting an inward movement towards the interhelical core in mPS-bound structures. The inward displacement was also captured by the decrease in the interhelical distances between TM6 and TM3. These results indicated that mPS binding shifted GPR174 from an active to an inactive conformational state. Comparative structural analysis with experimentally resolved inactive-state class A GPCRs further supported this conclusion (Fig. 3d).

Beyond global conformational changes, analysis of the shortest communication paths within GPR174 revealed differences in the allosteric signaling networks induced by LysoPS and mPS binding (Fig. 4a-b). In the LysoPS-bound state, extracellular interactions localized to TM6, TM5, and ECL2 were efficiently coupled to the conserved PIF motif, which was further connected to N/DPxxY and DRY motifs. These results are consistent with established models of class A GPCR activation, in which ligand-induced rearrangements are transmitted through conserved microswitches to the cytoplasmic G-protein–binding region^43,44^. In contrast, mPS binding redirected extracellular communication toward TM1 and TM4 but failed to establish productive connections to the PIF motif or other conserved elements of the receptor core. The absence of these links effectively decoupled the extracellular ligand-binding site from the intracellular signaling machinery, thereby preventing propagation of conformational changes necessary for G-protein activation. The communication pathway analysis aligns well with the distinct hydrogen binding profiles observed for LysoPS and mPS (Fig. 4c). LysoPS engages a broader set of residues with moderate and transient occupancy, reflecting its flexible pose and reduced stability within the pocket (Fig. 4e). In contrast, mPS forms a well-defined and persistent hydrogen bond network, most notably with R175^ECL2^ and residues on TM1, interactions absent in the LysoPS complex (Fig. 4f). These stable contacts likely lock the receptor in an inactive conformation and suggest that targeting TM1 and ECL2 interactions may be an effective strategy for designing GPR174 antagonists.

The conformational dynamics of LysoPS and mPS further illustrate how these ligands differentially stabilize GPR174. RMSD clustering analyses showed that mPS adopted a highly stable and dominant binding pose, whereas LysoPS displayed extensive conformational heterogeneity (Fig. 5a-b). Two key contributors to this heterogeneity were identified. One factor was the lateral displacement of the LysoPS head group from the pocket formed by TM1, TM2, TM3, and TM7 toward TM3, TM4, TM5, and TM6. This movement was also observed earlier in short simulations reported by Liang et al. (2023)^17^. Two possible explanations may account for this behavior. Structural modifications that enhance hydrophobic contacts within the receptor core are known to stabilize agonist binding. For instance, Ikubo et al. (2015)^8^ demonstrated that replacing the glycerol backbone of LysoPS with a meta-substituted benzene system improved agonist activity, likely by increasing vdW interactions. Our simulations supported this view, as mPS containing this chemical substitution showed more stable binding, stronger hydrophobic contacts, and reduced conformational heterogeneity compared to LysoPS. In addition, the absence of a bound G protein in our simulations may have facilitated conformational relaxation of the receptor–LysoPS complex. Biophysical studies have shown that agonist affinity is increased when the receptor is coupled to its G protein^45^. Thus, the lack of G-protein coupling in our simulations could have promoted destabilization of the LysoPS binding pose. This behavior may represent an intrinsic intermediate state of the GPR174-LysoPS complex following G-protein dissociation, preceding full receptor deactivation.

Another factor was the dynamic behavior of the LysoPS acyl chain, which transiently disengaged from the hydrophobic pocket between helices 4 and 5 to interact with surrounding lipids before re-entering the pocket (Fig. 5c-e). Together, these observations point to a potentially multi-step unbinding process in which membrane lipids can transiently occupy the hydrophobic pocket and compete with the LysoPS acyl chain, thereby modulating ligand dynamics. Such lipid competition highlights an active role of the membrane in shaping agonist binding stability and influencing intermediate receptor states. Consistent with this idea, lipid interaction analysis revealed distinct patterns of cholesterol and PIP2 binding in LysoPS-versus mPS-bound receptors (Fig. S7-S8). While both systems exhibited multiple cholesterol-binding sites, a site resolved in the experimental structure was only observed in the mPS-bound state (Fig. S9). PIP2 binding showed even stronger ligand dependence: the LysoPS-bound receptor accommodated a single PIP2 molecule between helices 4 and 5, overlapping with the displaced ligand tail, whereas the mPS-bound receptor revealed three distinct PIP2 binding sites that were absent in the agonist-bound complex (Fig. 6). These results suggest that PIP2 stabilizes distinct receptor conformations depending on ligand occupancy.

This study provides a comprehensive view of how LysoPS and mPS differentially modulate GPR174 dynamics, offering mechanistic insights into receptor activation and inhibition. mPS acts as a potent antagonist of GPR174 by promoting an inactive receptor conformation and stabilizing a well-defined hydrogen bond network, particularly through interactions with TM1 and ECL2. Unlike LysoPS, which displayed high conformational heterogeneity, mPS maintained a dominant and stable binding pose that effectively decoupled the extracellular ligand-binding site from the conserved motifs that allosterically relay activation signals to the G protein binding sites on the cytoplasmic end. The ability of mPS to restrict receptor flexibility and block communication to conserved activation motifs highlights its utility as a structural template for antagonist design. Beyond these ligand-specific effects, our results also highlight the broader role of lipid interactions in shaping receptor conformational dynamics and stabilizing distinct functional states. Collectively, these findings advance our understanding of GPCR functional plasticity and provide a framework for the rational design of selective GPR174 antagonists with potential immunotherapeutic applications.

## Methods

### Datasets

The structure of the GPR174 protein bound to the endogenous ligand LysoPS was obtained from the Protein Data Bank (PDB ID: 7XV3)^17^. The structure of a reported antagonist (compound 7d/mPS)^16^ of GPR174 was obtained with the help of an artificial intelligence-based platform (DECIMER.ai) that extracts chemical structures from scientific publications^46^.

### Molecular Docking

Molecular docking was performed to model an initial structure of GPR174 bound to mPS. The endogenous ligand LysoPS was also docked with similar input parameters to validate the docking protocol. The docked complex with LysoPS was aligned on a reference structure (PDB ID: 7XV3) to ensure accuracy of the docking method. Molecular docking was performed using PyRx (version 0.9.2), a graphical user interface for AutoDock Vina^47^. The GPR174 protein and ligand structures (LysoPS and Compound 7d) were imported into PyRx. The protein structure was converted to the AutoDock-compatible format (PDBQT) using the built-in PyRx tools, which added Gasteiger charges and defined the rotatable bonds. Similarly, the ligands were converted to PDBQT format after energy minimization by OpenBabel^48^. The docking grid was centered on the active site of GPR174. The grid box dimensions were set to encompass the entire binding pocket, allowing for sufficient space for the ligands to explore various conformations. The grid box was defined with the following parameters: center (x, y, z) coordinates [-36.07, 88.11, 10.03] and dimensions of [39.70, 46.20, 44.85] Å, ensuring coverage of the binding site and surrounding regions. For each ligand, a total of nine poses were generated, and the conformations were ranked based on their binding affinities. The exhaustiveness parameter was set to 8 to balance computational efficiency and thoroughness of the search. The docking results were analyzed to identify the binding poses with the lowest binding free energy, and these were selected as the most favorable conformations.

### Modeling membrane-bound GPR174 complexes

The crystal structure of GPR174-LysoPS (PDB ID: 7XV3) and the docked GPR174-mPS were placed in a heterogeneous lipid bilayer using the CHARMM-GUI server^49^. The N- and C-termini of GPR174 were patched with NTER (NH3+ group) and CTER (COO-group) to reflect the common protonation states at neutral pH. The disulfide bond between Cys91 and Cys168 was preserved in both complexes. Consistent with previous literature on protonation states in active state conformations of GPCRs^17,50^, the aspartates at positions 65 and 16 were protonated in the LysoPS-bound GPR174 and both these residues were kept unprotonated in the mPS-bound complex. We utilized the Orientations of Proteins in Membranes (OPM) database^51^ to orient the protein inside the lipid bilayer (Fig. 1c) composed of Chl, PSM, PC, PE, Pla(PE), PS, and PIP_2_ ^20,52^. The systems were then solvated using the TIP3P water model. To neutralize the systems and maintain an ionic strength of 0.15M, sodium (Na⁺) and chloride (Cl⁻) ions were added. Finally, the simulation input files were prepared for GROMACS setup ^53^ using the CHARMM36m force field, setting the temperature to 310K under the NVT (constant number of atoms, volume and temperature) ensemble.

### Simulation Setup

The GROMACS-compatible outputs from the CHARMM-GUI server were subjected to energy minimizations followed by multi-step equilibrations. Minimization was carried out using the steepest descent integrator until the maximum force converged to below 1000 kJ/mol/nm^2^. Harmonic position restraints were applied to protein backbone atoms and ligand heavy atoms with a force constant of 4000kJ/mol/nm^2^. Protein side-chain atoms and lipid phosphorus atoms were restrained with force constants of 2000 kJ/mol/nm^2^ and 1000kJ/mol/nm^2^, respectively. Additionally, dihedral restraints with a force constant of 1000kJ/mol/rad^2^ were applied to different sets of atoms to maintain specific conformations of the lipid molecules. Afterwards, the system was equilibrated in a stepwise manner to gradually relax the atomic positions. This process involved six successive NVT runs, each with decreasing force constants. In the final equilibration step, the restraints on the protein side chains and lipids were completely removed, leaving only minimal restraints on the protein backbone and ligand heavy atoms, which were also eventually removed for the final production run. Production runs were performed in duplicates for both systems, each initiated with different randomized initial velocities. All four systems were subjected to an individual MD run of 2 µs using the leap-frog integrator with a 2 fs time step. One of the LysoPS-GPR174 runs was extended to 4 µs. The simulations were performed under periodic boundary conditions using a Verlet cutoff scheme with neighbor lists updated every 20 steps. The short-range electrostatic and van der Waals interactions were both set with a cutoff of 1.2 nm. Long-range electrostatic interactions were handled using the Particle Mesh Ewald (PME) method^54^ with all covalent bonds involving hydrogen atoms constrained using the LINCS algorithm^55^. The temperature was maintained at 310 K using a Nose-Hoover thermostat. The temperature coupling was applied semi-isotropically to the solute (protein-ligand), membrane, and solvent (water and ions) groups, each with a time constant of 1.0 ps. The pressure was set to 1.0 bar using a Parrinello-Rahman barostat^56^ with a time constant of 5.0 ps. Coordinates, velocities, and energies were written out every 10 ps.

### Trajectory Analysis

The trajectories were analyzed mostly with GROMACS tools and MDAnalysis library^57^. Cα atoms RMSD and RMSF were computed using GROMACS analysis tools. Bilayer thickness (P-P distance) was calculated as the z-distance between the center-of-mass of phosphorus atoms in the upper and lower leaflets. Interhelical angles were calculated with *gmx gangle*, using vectors defined by selected Cα atoms of each helix. The *gmx dist* tool was employed to measure the cytosolic-end distance between transmembrane helices TM6 and TM3, computed as the distance between the centers of mass of five selected Cα atoms on each helix. Conformational clustering was performed using the *gmx cluster* tool with the GROMOS algorithm^58^ with an RMSD cutoff of 0.18 nm. Hydrogen bond analysis included both the number and occupancy of hydrogen bonds, calculated using *gmx hbond* and the plot_hbmap_generic.pl script, respectively. Protein structure network (PSN) analysis was conducted using PSNtools^24^, with a frequency cutoff of 60%, such that only links and hubs present in at least 60% of the trajectory frames were considered. Node size and color were mapped to interaction force and frequency, respectively. For protein-lipid interaction analysis, PyLipID^30^ was used with modifications to consider only heavy atoms of the protein. For lipid molecules, all heavy atoms were included for Chl, while for PIP2 the acyl chains were excluded. A 0.4 nm distance cutoff was applied to define lipid–protein interactions. Visualizations were produced using PyMOL and data plots were generated using Grace.

## Acknowledgements

All simulations were run on Lonestar6, a high-performance computing system provided by the Texas Advanced Computing Center. AAG acknowledges financial support from the National Institutes of Health Institute of General Medicine grant R01GM144836. VKB was supported in part by a Cancer Therapeutics Training Program fellowship (CPRIT Grant No. RP210043).

