## Supplementary Data for "GPR174 Antagonism: Structure, Function, and Dynamics"

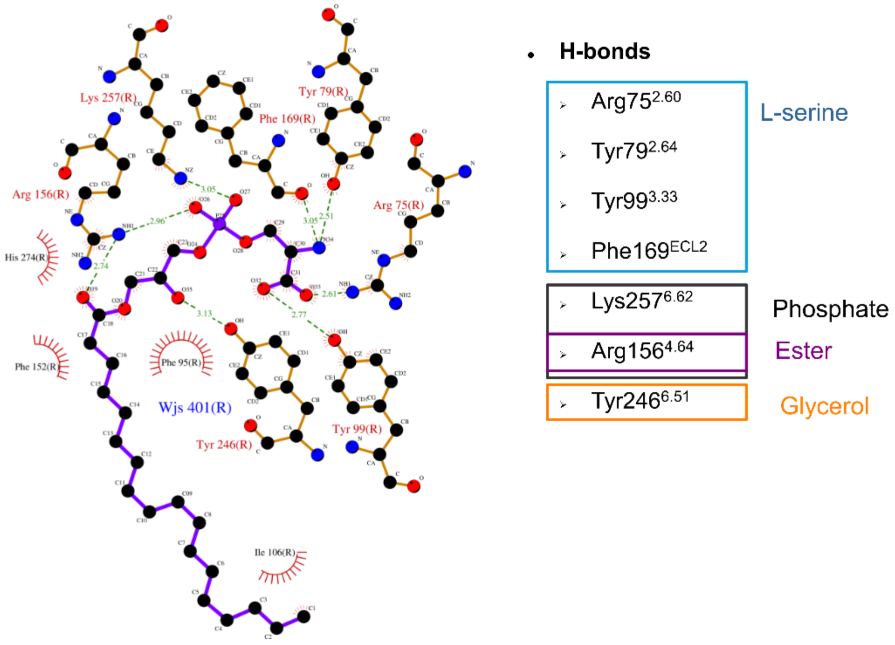


Fig. S1. **Two-dimensional interaction map of LysoPS bound to GPR174**, derived from the experimental structure. Interacting residues listed inside colored boxes correspond to different LysoPS chemical groups.

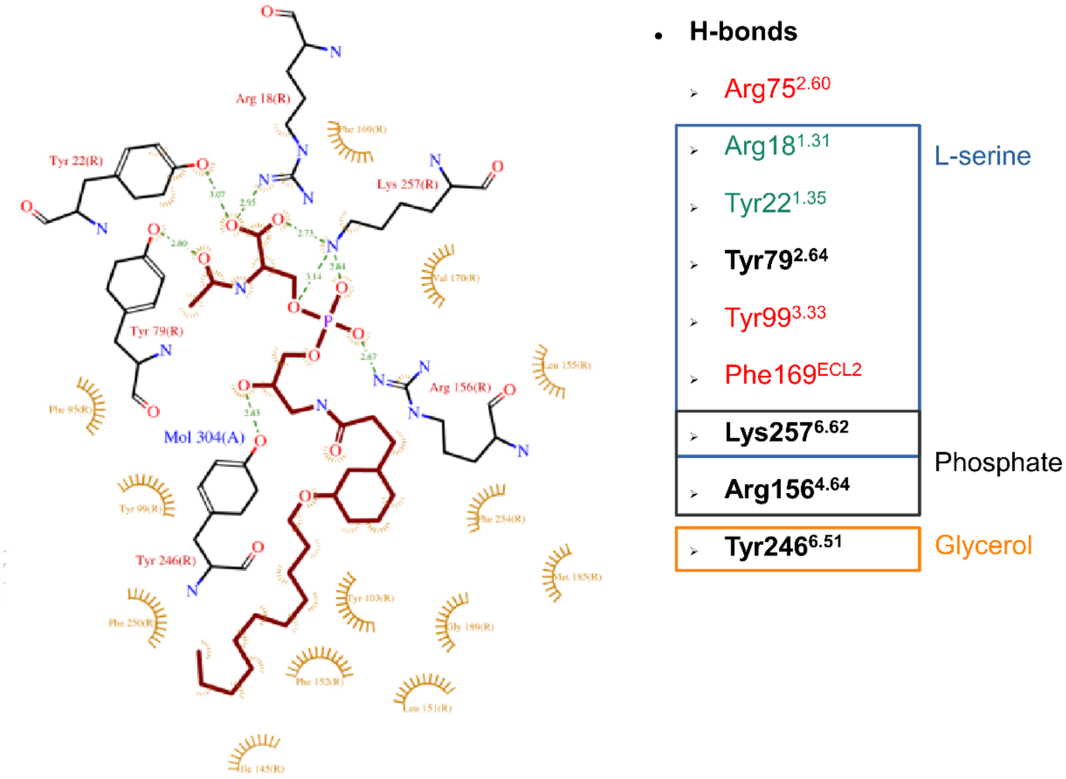


Fig. S2. **Two-dimensional interaction map of docked mPS with GPR174**. Residues are colored according to differences relative to the LysoPS binding pose: red (lost interactions), green (novel interactions), and black (conserved interactions). Interacting residues listed inside colored boxes correspond to different mPS chemical groups.

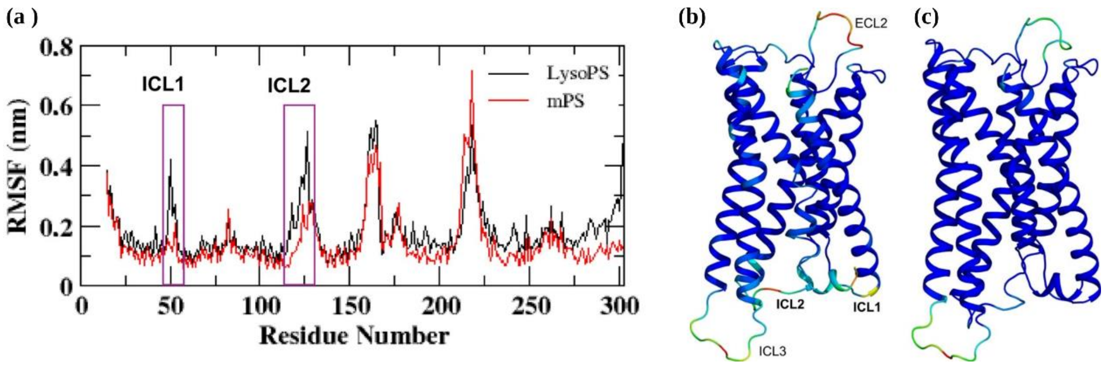


Fig. S3. **Analysis of fluctuations at the residual level**. (a) Analysis of root mean square fluctuation (RMSF) of backbone atoms for the GPR174 protein bound to LysoPS (black) or mPS (red). The regions corresponding to intracellular loops 1 (ICL1) and 2 (ICL2) are highlighted with purple boxes. Cartoon representations of the average structures of GPR174 bound to (b) LysoPS and (c) mPS with RMSF values mapped onto the protein as B-factors. The color scale ranges from blue (low flexibility) to red (high flexibility).

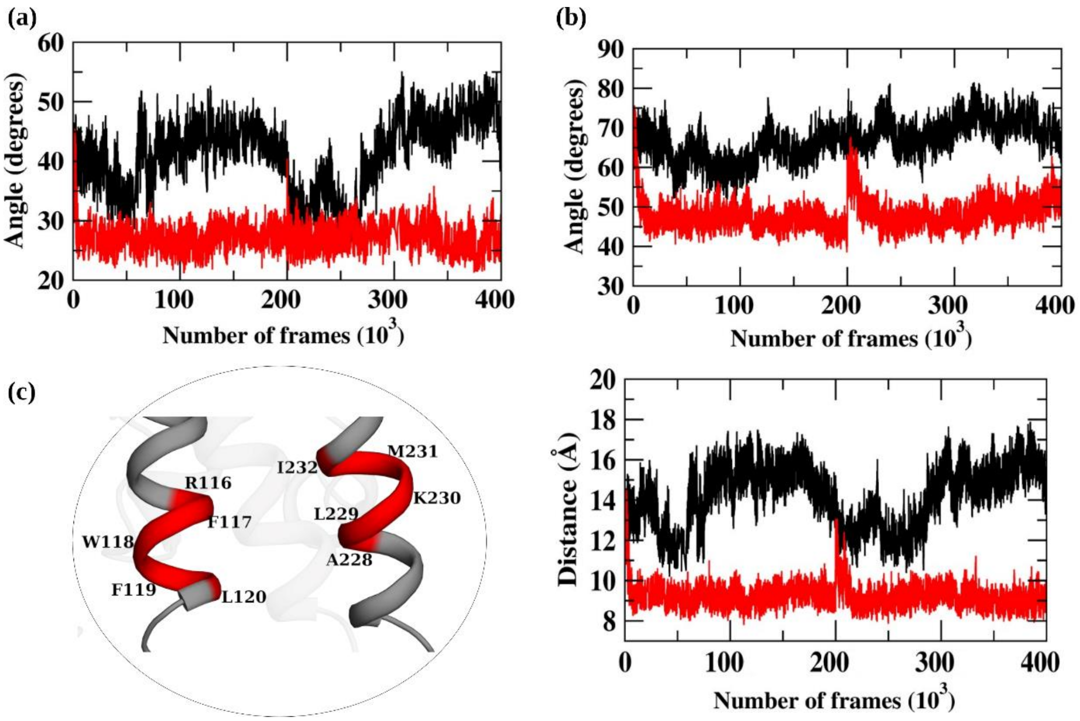


Fig. S4. **mPS binding inactivates GPR174**. Time evolution of the angle formed between (a) TM6 and TM3 and (b) TM6 and TM2 at the cytoplasmic side of GPR174 bound to LysoPS (black) and mPS (red). (c) Structural representation highlighting residues for calculating the distance between TM3 and TM6. The corresponding time evolution of distances calculated between the center-of-mass of selected atoms is shown on the right.

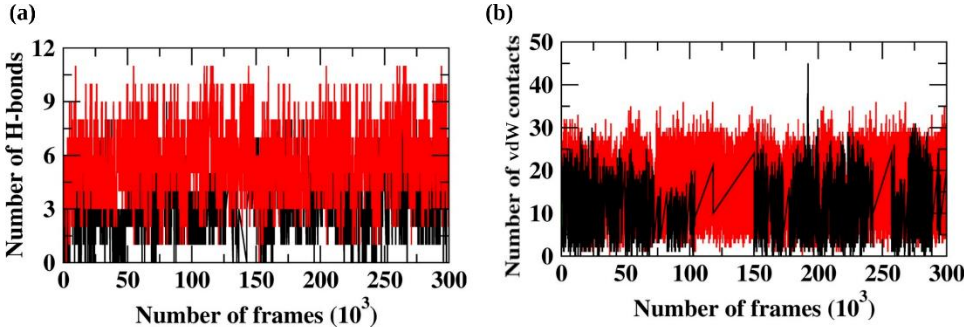


Fig. S5. **Analysis of non-bonded interactions**. Time evolution of (a) number of hydrogen bonds and (b) number of van der Waals contacts formed by LysoPS (black) and mPS (red) with GPR174.

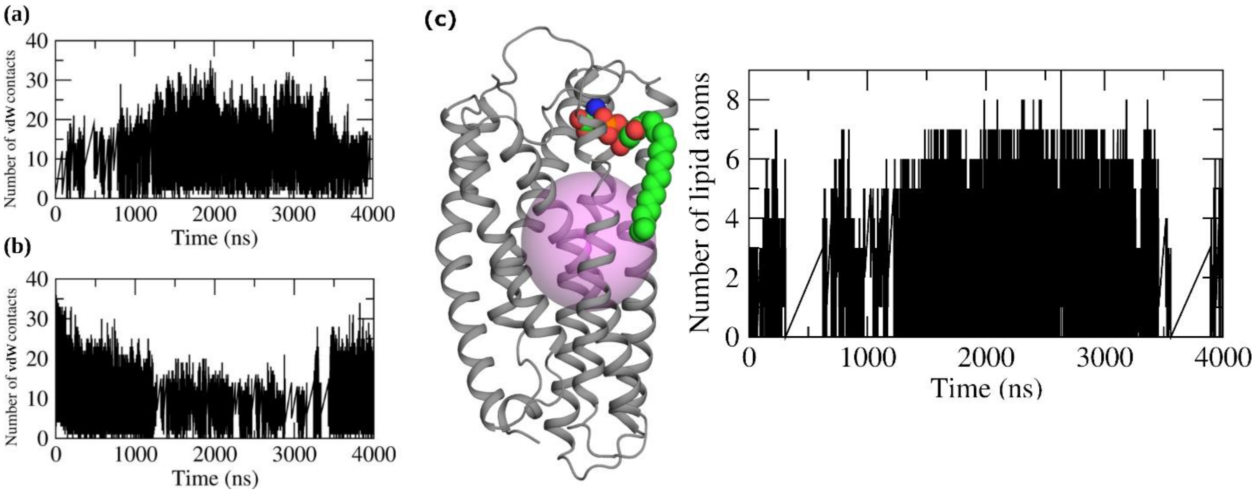


Fig. S6. **Analysis of LysoPS acyl chain binding dynamics**. Number of van der Waals (vdW) contacts formed by LysoPS acyl chain with (a) membrane lipids and (b) protein. All side chain C atoms of hydrophobic residues (Ala, Val, Iso, Leu, Met, Phe, Tyr, and Trp) of GPR174, C atoms of LysoPS acyl chain, and C atoms of acyl chains of membrane lipids within 4Å of distances were considered to form vdW contacts. (c) A representative snapshot of the GPR174 receptor with a defined pocket volume (magenta sphere) used to quantify membrane lipid occupancy. The accompanying plot to the right displays the number of membrane lipid atoms within this specific pocket as a function of simulation time.

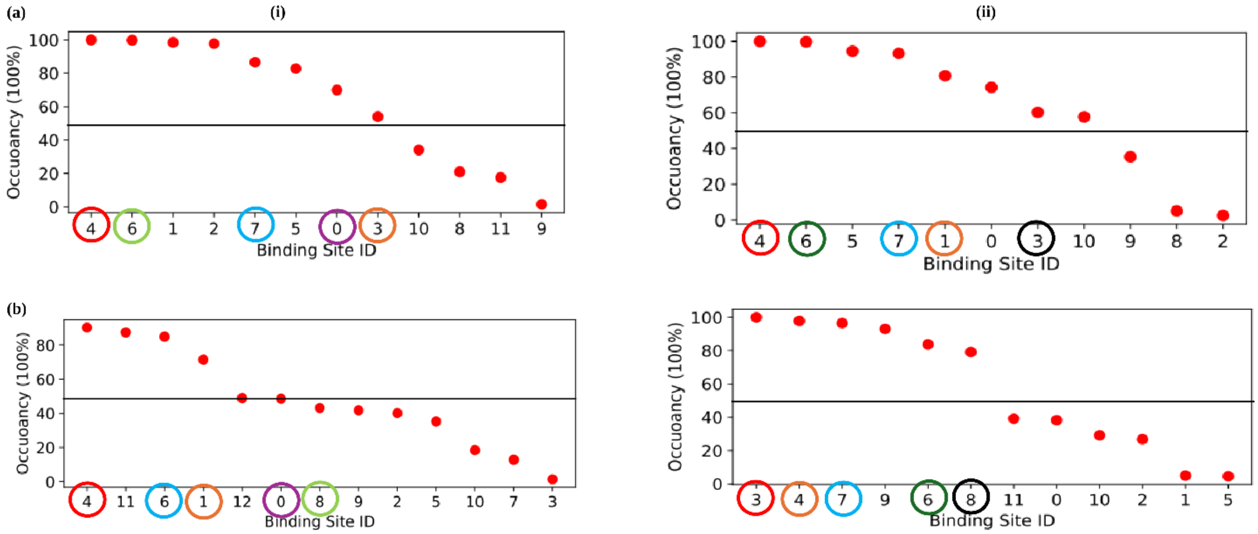


Fig. S7. **Chl-GPR174 interactions**. The occupancy of cholesterol binding sites on the GPR174 receptor in (i) LysoPS and (ii) mPS during two independent molecular dynamics simulation runs (a: Run 1; b: Run 2). The horizontal black line indicates a 50% occupancy threshold. The horizontal black line marks the 50% occupancy threshold. Binding sites that are conserved across both runs are highlighted with the identically-colored circles for comparison.

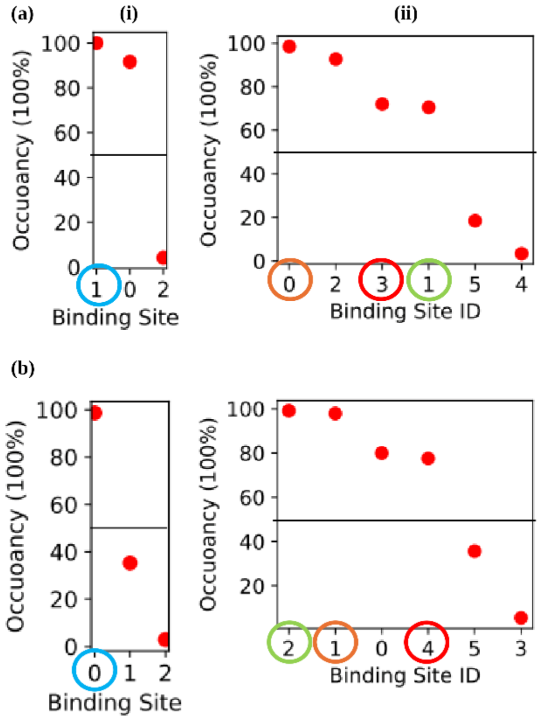


Fig. S8. **PIP$_2$-GPR174 interactions**. The occupancy of PIP2 binding sites on the GPR174 receptor in (i) LysoPS and (ii) mPS during two independent molecular dynamics simulation runs (a: Run 1; b: Run 2). The horizontal black line indicates a 50% occupancy threshold. The horizontal black line marks the 50% occupancy threshold. Binding sites that are conserved across both runs are highlighted with the identically-colored circles for comparison.

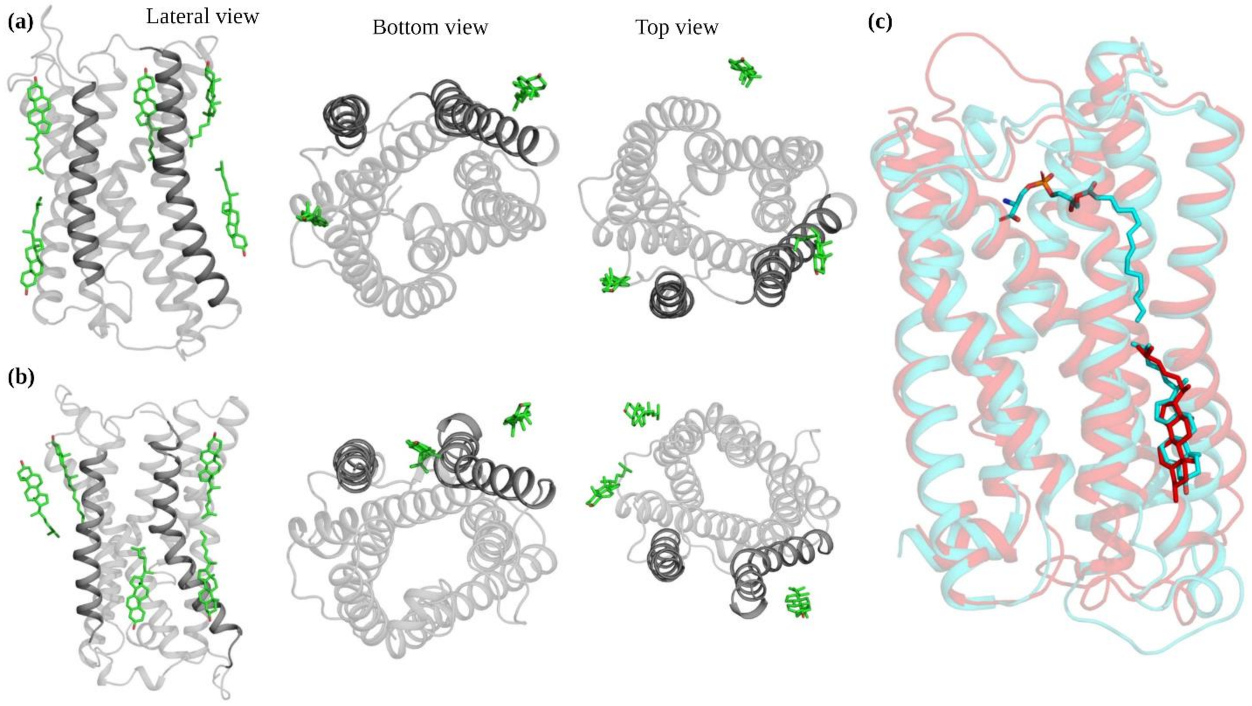


Fig. S9. **Protein-lipid interactions**. Analysis of cholesterol binding sites for (a) LysoPS-bound and (b) mPS-bound GPR174 shown in lateral, bottom, and top views. Cholesterol molecules are depicted in green sticks, and the receptor is shown in cartoon representation with helices (TM4 and TM5) highlighted. (b) Superimposition of the conserved cholesterol binding site obtained from our analysis (red) on the experimental structure (PDB: 8KH5) of GPR174 (cyan), confirming overlap between predicted and experimental sites.
